# An Evaluation of Curriculum change in Physiology: A Mixed method research design

**DOI:** 10.1101/2021.04.19.440394

**Authors:** Shivayogappa. S. Teli, Senthil velou M., Soundariya K., Deepika Velusami, Senthamil selvi K, Mangani Mangalavalli S

**Affiliations:** Department of Physiology, Sri Manakula Vinayagar Medical College and Hospital, Puducherry; Department of Physiology, AIIMS Mangalagiri, Andhra Pradesh

**Keywords:** Physiology, Case-based learning, Likert scale, teamwork, communication skills, self-directed learning, problem-solving ability

## Abstract

**Introduction:** The quality of both teaching and learning techniques in health sciences determines the competency of the doctors produced and the patient care. Realizing the necessity of active learning at the undergraduate level, curricular reforms were crucial to ensure that students play an active role in the learning process and imbibe all the prerequisite qualities of a competent health professional. The objective of this study was to implement and evaluate case-based learning in the physiology curriculum.

**Methods:** The study included 150 first-year MBBS students. We followed a mixed research design in this study. A short lecture on anemia was followed by two sessions of CBL with a gap of one week. A structured questionnaire on a 5-point Likert scale was used to collect students’ perceptions. The internal consistency of the questionnaire was Cronbach α-statistic = 0.8. Faculties’ feedback was collected through Focus Group Discussion (FGD).

**Results:** Out of 145 participants, 117 responded to feedback. Students perceived that the CBL method was effective in understanding anemia topic (71%); promoted meaningful learning (83%); encouraged teamwork (69%); improved communication skills (65%); helps in future application of knowledge (81%); motivated self-directed learning (66%); helped to understand physiology concepts better (72%); leads to the development of problem-solving abilities (70%); and better student-teacher relationship (72%). Faculties suggested developing an assessment plan for future CBL sessions.

**Conclusion:** In our experience, CBL is an effective, active teaching-learning tool that improves students’ understanding of basic concepts, clinical knowledge, problem-solving abilities, teamwork, communication skills, student-teacher relationship, and self-directed learning.

## Introduction

“Progress is impossible without change, and those who can’t change their minds can’t change anything”-George Bernard Shaw.

The journey of life itself, from conception to culmination, is the best example to – Change. The process of education - acquiring knowledge, skills, values, morals and the way how it is acquired has also seen profound changes over the centuries in its structure, understanding, and delivery. The purpose of Medical education in India or in any other country is to train skilled health professionals who would serve society diligently to prevent, cure, and promote the well-being of mankind [1]. The quality of both teaching and learning techniques in the health sciences determines the competency of the doctors produced and the patient care.

The majority of medical colleges in India teach basic sciences in 1st-year MBBS by traditional methods alone, where the curriculum decided by universities and higher governing bodies will be delivered to the students via lectures and practical sessions [1,2]. The traditional style of teaching isn’t well-received by the new generation of students who feel it as boring, less interactive, teacher-oriented, and sleep-inducing. Such a method of teaching and learning is teacher-centered with minimal active participation from the students. There is a widespread belief among medical educators throughout the world that lecture-based teaching alone is insufficient to address the needs of all learners and is not suitable for teaching higher-order cognitive skills, such as synthesis, analysis, and application, which are critical for medical practitioners [3]. When the students’ involvement is less in the learning process, obviously the outcome will also be less productive.

Realizing the compelling necessity of active learning at the undergraduate level, curricular reforms were crucial to ensure that students play an active role in the learning process and imbibe all the prerequisite qualities of a competent health professional [1]. Physiology is one of the basic sciences in 1st MBBS and needs to be taught and learned effectively so as to be placed in the context of disease when the medical students start attending their clinical postings during 2nd MBBS. The emerging trend all over the world is to have a problem-based, integrated, student-centered medical curriculum that allows active participation from the students and facilitates self-directed learning [4].

There are many different ways of explaining how adults learn effectively [5]. Likewise, there are many different learning strategies that allow active participation of learners and the development of better knowledge, skills, critical thinking, and values [6]. Case-based learning (CBL) is a long-established pedagogical method where basic sciences’ concepts are studied in relation to clinical conditions. CBL enjoys a variety of definitions across various disciplines and contexts. Basically, it is a form of inquiry-based learning that uses clinical cases to aid teaching. It fits on the continuum between structured and guided learning [7,8]. CBL integrates preclinical and clinical subjects creating a link between theory and practice allowing students to think holistically about the profession. The advantages of case-based learning are manifold: promotion of self-directed learning; development of clinical reasoning; acquisition of clinical problem solving, decision making, and communication skills; and stimulation of deep learning [3,4,7-12]. Although many countries have shifted their curriculum from teacher-centered to student-centered training, the majority of medical colleges in India are still practicing old-fashioned teaching. On literature search, we found that very few Indian authors have reported the inclusion of active learning methods like CBL in their curriculum [3,4,10,11].

Although traditional teaching is useful in covering a vast amount of information to larger audiences, they are not effective for active learning. The learners remain passive and less attentive in the classroom leading to reduced perception of information. In our institution also, physiology was being taught mainly by traditional style to convey the basic and applied concepts of health and diseases. It is the right time to change our teaching styles and embrace modern, active teaching-learning methods in our curriculum.

There were a few reasons why we wanted to initiate the changes in our existing teaching styles and the curriculum. Firstly, it was the training of our faculties in the faculty development program (FDP), which changed our awareness of teaching and learning in medical education. Secondly, it’s the students’ feedback on learning physiology. Students felt that learning physiology was boring and difficult. Many physiology concepts were imaginary and complex that were difficult to learn and understand. And lastly, yet the essential one, it’s the feedback from our colleagues in the clinical departments who expressed their concerns over students’ acquisition of prerequisite knowledge and skills before their first clinical postings.

The primary objective of this educational research was to design and implement case-based learning in our curriculum to facilitate active learning among the students. In addition, the secondary objective was to evaluate the stakeholders’ perception for improvement and future implementation of CBL.

## Materials and Methods

This study was conducted in the Department of Physiology of Sri Manakula Vinayagar Medical College and Hospital, Puducherry. Institutional Research and Ethics committee permissions were taken. The study involved all 150 students of first-year MBBS (2017-18 batch). We used a mixed research design (both quantitative and qualitative) in the current study. The framework of the study is outlined below.

### Framework of CBL

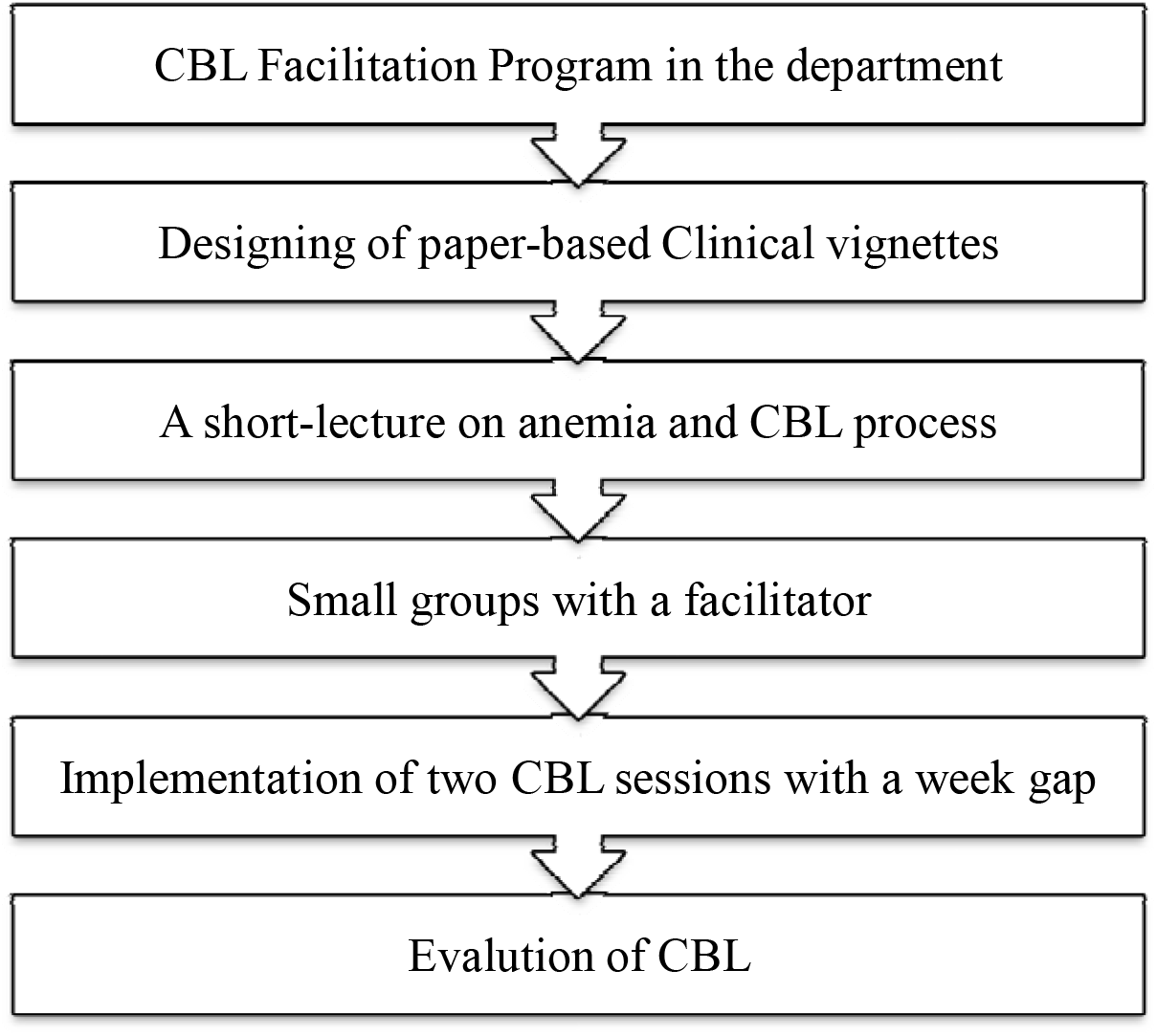

To this end, a department meeting was arranged with all the faculties, to solicit their cooperation in the study. In the meeting, all concurred with two decisions: firstly, to include ‘anemia’ topic to teach in CBL and secondly, all faculties must be trained before the implementation of CBL. The principal investigator took the responsibility to arrange a ‘faculty training’ session and to prepare anemia-related clinical scenarios.

Later, a ‘CBL Facilitation Program’ was arranged in the department with the help of the Medical Education Unit (MEU). It was truly an interactive session, which provided us an opportunity to understand the purpose of CBL, the role of teachers in CBL, how to prepare and validate paper-based clinical scenarios, and how to collect feedback from all the stakeholders. Two clinical scenarios – Microcytic and Macrocytic Anemia – were prepared and validated in the next 15-days period.

It was also felt necessary that students must be briefed about new learning intervention as it completely differs from traditional learning. The principal investigator explained to the students in brief about the CBL - its design, purpose, and role of students and teachers in the learning process. It was also emphasized that students must keep a note of how CBL influences their learning and give honest feedback when asked.

One of the important characteristics of CBL, which the CBL-proponents believe, is that the learners should have some background knowledge on the chosen topic. Therefore, a didactic lecture was taken before CBL intervention on erythrocyte structure, function, production, factors regulating erythropoiesis, and a brief introduction to anemia.

We planned to conduct two CBL sessions with a gap of one week. Case scenarios on Microcytic and Macrocytic Anaemia were selected for the first and second sessions respectively. We usually have tutorial sessions every Friday between 2.30–4.30 pm. The idea of giving one week gap in between two learning sessions was to provide sufficient time for self-directed learning (SDL). Whatever additional information they gain during this period would definitely help them in solving the new clinical vignette given during the next CBL session. Moreover, it helps the teachers to know whether SDL is actually happening or not.

As a start-up to implement CBL, all 150 students were divided into 5 tutorial groups and each group had a trained faculty to facilitate the session. Each tutorial group (30) was again subdivided into three small groups with 10 students each to enhance active involvement and learning from all.

The process of CBL during these two sessions was similar. Each CBL session was given two hours time to complete: initial 10 minutes to read and understand about the given clinical scenario; next 60 minutes to discuss and solve all the specific learning objectives (SLOs) by self-learning and group discussion; another 30 minutes to interact with teachers to get guidance; and last 20 minutes for final conclusion. One of the students from each group, who voluntarily came forward, was given the opportunity to read the scenarios to the rest of the group. Students were also allowed to take a snapshot of the case on their mobile and read for themselves. They referred to textbooks to find the solutions for SLOs during self-learning and group discussion. It is worth mentioning here that the whole purpose of CBL is to encourage learners towards active learning. Teachers’ role was to give them guidance wherever students deviated from the main topic, ensure that they actively participate in the learning, and achieve all SLOs by end of the session. During the second CBL session, which happened next Friday, a new clinical scenario was given to all the groups to discuss and solve the related SLOs. The CBL method proposes that learners acquire new information by self-learning and become more competent in problem-solving with regular exposure to CBL.

### Data collection and analysis

Students’ feedback was collected after completing both the CBL sessions. It was an anonymous survey and private identities like name, roll number, age, and gender were kept confidential. A structured questionnaire on 11 different items (understanding, teamwork, communication skills, SDL, student-teacher relationship, problem-solving abilities, future application, etc) was used to collect students’ perception on CBL. The internal consistency of the questionnaire was measured using the Cronbach α-statistic (Cronbach α-statistic = 0. 8). Out of 11 questions, ten questions were closed type, which required rating on 5-point Likert scale (indicating students’ level of agreement from strongly agree to strongly disagree) and the last question was an open-ended type [26]. As it was a program evaluation (Kirkpatrick level 1), the items were aligned with the purpose of CBL. Faculties’ feedback was collected through Focus Group Discussion (FGD). For quantitative analysis, simple frequency distribution table was used to summarize the data and a simple descriptive narrative was preferred for qualitative analysis of open-ended question and FGD data [27].

## Results

Out of 150 students, 145 participated in both the CBL sessions and 117 responded to written feedback (response rate was 80.68%). All the faculties (n=7) gave their feedback during Focus Group Discussion.

### Students’ feedback

The results of the students’ responses to CBL questionnaire (Table 1) revealed that the majority of students agreed or strongly agreed that CBL method was more effective in understanding anaemia topic (83 of 117 students, 71%); promoted meaningful learning (97 of 117 students, 83%); encouraged team work (81 of 117 students, 69%); improved communication skills (76 of 117 students, 65%); helps in future application of knowledge (95 of 117 students, 81%); motivated self-directed learning (82 of 117 students, 66%); helped to understand physiology concepts better (84 of 117 students, 72%); leads to development of problem-solving abilities (82 of 117 students, 70%); and better student-teacher relationship (83 of 117 students, 72%).

**Table 1.** shows Students’ responses to CBL questionnaire.

| S.No | Likert items/statements | Likert response options |  |  |  |  |
| --- | --- | --- | --- | --- | --- | --- |
|  |  | SA*<br>n(%) | A*<br>n(%) | NAN<br>D*<br>n(%) | D*<br>n(%) | SD*<br>n(%) |
| 1 | In understanding anaemia topic, CBL sessions were useful | 33(28) | 50(43) | 31(26) | 2(2) | 1(1) |
| 2 | Relevant and Interesting case scenarios were given to learn anaemia | 37(32) | 47(40) | 28(24) | 5(4) | 0(0) |
| 3 | CBL promotes meaningful learning than the regular lecture classes | 46(39) | 51(44) | 14(12) | 5(4) | 1(1) |
| 4 | CBL encouraged me to work with my friends as a "team" in solving case scenarios | 46(39) | 35(30) | 25(21) | 8(7) | 3(3) |
| 5 | CBL provides an opportunity to interact with friends and improve communication skills | 40(34) | 36(31) | 32(27) | 7(6) | 2(2) |
| 6 | CBL method is useful in the future application of knowledge | 54(46) | 41(35) | 21(18) | 1(1) | 0(0) |
| 7 | CBL motivated me to refer more learning resources like textbooks, online search, etc. (self-directed learning) | 30(26) | 52(44) | 30(26) | 5(4) | 0(0) |
| 8 | CBL facilitates a better and healthy student-teacher relationship | 35(31) | 48(41) | 26(22) | 4(3) | 4(3) |
| 9 | CBL helped me to develop problem-solving abilities | 30(26) | 52(44) | 30(26) | 5(4) | 0(0) |
| 10 | CBL improved my understanding of physiology concepts better | 34(29) | 50(43) | 28(24) | 4(3) | 1(1) |
| 11 | Briefly mention your learning experiences/comments/suggestions related to CBL sessions |  |  |  |  |  |

\*SA=Strongly Agree, A=Agree, NAND=Neither Agree Nor Disagree, D=Disagree, and SD=Strongly Disagree. n=number of students

In the open-ended question, students were requested to share suggestions/comments/experiences on CBL sessions (refer Table 2). Few comments on CBL are as follows:

**Table 2.** shows Students’ responses to open-ended questions in CBL.

| S. No | Suggestions/Comments/Experiences | Percentage |
| --- | --- | --- |
| 1 | To conduct more CBL sessions on clinically important topics | 60 |
| 2 | To reduce number of participants in each group | 46 |
| 3 | Helps in peer learning | 42 |
| 4 | Learn diagnosis skills | 34 |
| 5 | Acquire clinical knowledge from many other sources | 39 |
| 6 | Freedom to learn in our own way | 25 |
| 7 | Improves relationships among students | 21 |
| 8 | To give assignments on the CBL topics | 10 |
| 9 | Enjoyed new method of learning | 30 |

- “Learned diagnosis skills”
- “Freedom to learn”
- “Acquired clinical knowledge”
- “Referred additional books”
- “Enjoyed new method”
- “Improves relationships with friends”

They also suggested conducting more CBL sessions in the future, and to give assignments on clinically important topics so that they can study it later. Students’ suggestions requesting for more CBL sessions and assignments, indicate that they thoroughly enjoyed CBL method and become motivated to learn.

### Faculties’ feedback

Faculties’ feedback was collected on the following parameters: their experience with CBL sessions in comparison to regular teaching; their perception on students’ participation and learning in CBL (refer Table 3). All teachers expressed that CBL gave them a different teaching experience altogether. They were also surprised by students’ active involvement in CBL discussion. However, they pointed out two important areas of concern related to CBL sessions. Firstly, there should be less number students in each group so that monitoring and facilitation becomes easy. Secondly, they all felt that we must develop a plan to assess what students have learnt in CBL session. It’s true; there should be a measure to know the progress of learning. And, all the health educators very well know, “Assessment drives learning.”

**Table 3.** shows faculties’ responses to CBL (n=7).

| S. No | Experiences/Comments/Suggestions |
| --- | --- |
| 1 | It's a different teaching experience |
| 2 | CBL session covered only small topics |
| 3 | Good for teaching clinical related topics |
| 4 | Learning through CBL sessions should be assessed |
| 5 | Facilitation and monitoring was difficult |
| 6 | Students enjoyed entire session |
| 7 | Students were active and didn't sleep |

## Discussion

The purpose of medical education in India is to create competent health professionals who recognize “health for all” as a national goal. It is the collective responsibility of the Medical Council of India (now, National Medical Commission), medical colleges, and health educators in India to train the skilled Indian Medical Graduates (IMGs) who can effectively provide preventive, promotive, curative, palliative, and holistic care to the patients they serve [1]. It can be successfully achieved only by step-wise efforts beginning from pre-clinical years to internship level.

The objectives of this study were to design and implement CBL in the physiology curriculum to encourage active learning and also to evaluate stakeholders’ perceptions of CBL implementation. The overall results indicate that students showed a strong preference for learning via CBL as it facilitates a better understanding of physiology concepts, meaningful learning, and future application of knowledge. Learning Physiology through systems-based didactic lectures alone is difficult for beginners, as it largely includes new medical terminologies, imaginary concepts, and less interaction with teachers [21]. Guided learning, like in CBL, facilitates effective learning of basic sciences, which in turn is essential for better understanding of clinical subjects, interpretation of patients’ clinical signs and symptoms, and analysis of laboratory results [16-21]. This kind of exposure and training is essential for the learners to excel in clinical years.

In addition, the majority of the students felt that CBL encourages teamwork and the development of communication skills, as it allowed them to interact, discuss and share with each other during the learning process. Teamwork in the health care system, where two or more people interact and work for a common purpose and goal, is a key component to make decisions as a unit, while giving patient care and communicating with patients’ family and friends. Undoubtedly, repeated CBL exposure in the pre-clinical years, prepares these budding doctors for their future responsibility of handling the situations in a real context [11,13-15,18-20].

Moreover, they felt that this kind of learning helps to acquire problem-solving ability. The problem-solving ability, which involves mental processes to solve medical problems, can be defined as a hypothetical-deductive activity engaged in by experienced physicians, in which the early generation of hypotheses influences the subsequent gathering of information [22]. Health professionals develop this virtue gradually over the years by accumulation of medical knowledge and experience. Learners can be trained to acquire such skills early in their profession by exposing them to clinical case discussion and decision-making processes. CBL is one of such techniques to train the problem-solving ability of future doctors and the quality of their decisions in patient care.

The educational environment in any discipline will have a definite impact on how students learn and progress. Amongst many factors, curriculum, teaching-learning techniques, type of teachers and their teaching styles, and student support system are few that decide the learning environment, student’s satisfaction, and academic achievement [23]. A friendly, supportive learning environment contributes to student well-being and enhances student empathy, professionalism, and academic success [24]. In our research, students agreed that CBL provides such a learning environment and strengthens the student-teacher relationship.

Current research also observed that CBL stimulates self-directed learning (SDL) among learners. Students accepted in the feedback that they referred more learning resources during and also in-between the two CBL sessions to solve the cases provided. Nowadays, SDL has gained more attention and popularity among health educators all over the world. In simple terms, it is a learning process in which learners take the initiative and responsibility for their own learning. One way of inculcating SDL among students is to give them case-based scenarios and guide the learners with questions, leading them to answers using recommended learning resources [25]. It enables health professionals to continue learning and updating their knowledge.

To the open-ended question, students responded by saying that more CBL sessions should be conducted on clinically important topics and suggested reducing the number of participants in each group. Students enjoyed learning in CBL sessions as it provides an opportunity to interact with each other, share their findings, make decisions, and more importantly they have the freedom to learn at their own pace. Learning any new information requires interest and involvement. CBL creates interest in learners’ minds by exposing them to new clinical scenarios and showing them the links between theory and practice.

It is worth mentioning here that successful implementation of any new curriculum requires adequate training and cooperation of faculties. During the focus group discussion, faculties shared their experience on the CBL method. They all felt that it was a different teaching experience compared to regular classroom teaching. The role of teachers in CBL is just to facilitate the learning process by guiding the students appropriately towards solving learning objectives. Our faculties mentioned that they can become better facilitators only after repeated participation in CBL. They felt that the major concern of the CBL method is that in a two-hour session, only small topics can be covered and it is a good method for teaching clinically related topics, but not for all. Surprisingly, everyone shared that case-based learning is an effective teaching method as all students were active throughout the session and didn’t sleep at all.

### The way forward

In our analysis, we identified few areas of improvement related to CBL implementation. Firstly, we need to select only the clinically important topics in physiology for teaching CBL. Secondly, an assessment for learning should be planned for CBL sessions. Thirdly, a decision has to be taken on how to conduct more CBL sessions in small groups with a minimum number of participants in each group. And finally, we have to work out giving assignments on CBL sessions to facilitate self-directed learning. The findings of the study were shared with all the faculties in the department and an evaluation report along with the future plan of action was submitted to the institutional curriculum committee for review and suggestions.

## Conclusion

Medical education is a dynamic process that keeps changing with the advent of new information and innovative teaching-learning methods. Our aim was to facilitate the learning process by implementing CBL in physiology. In our experience, CBL is an effective, active teaching-learning tool that improves students’ understanding of basic concepts, clinical knowledge, problem-solving abilities, teamwork, communication skills, student-teacher relationship, and self-directed learning. So, we need to adapt to effective learning strategies to train our students better as we all know, “Today’s learners will be tomorrow’s doctors.” Embracing new ideas and developments in teaching, changes our concept of training and learning in medical education. These little changes made in the curriculum, opened new opportunities for us to a novel learning experience.

## Acknowledgement

We heartily express our thankfulness to all the students of 2017-18 batch of Sri Manakula Vinayagar Medical College and Hospital for their active participation and feedback.

